# Multiomic clocks to predict phenotypic age in mice

**DOI:** 10.1101/2025.04.30.651114

**Authors:** Daniel L. Vera, Patrick T. Griffin, David Leigh, Jason Kras, Enrique Ramos, Isaac Bishof, Anderson Butler, Karolina Chwalek, David S. Vogel, Alice E. Kane, David A. Sinclair

## Abstract

Biological age refers to a person’s overall health in aging, as distinct from their chronological age. Diverse measures of biological age, referred to as “clocks”, have been developed in recent years and enable risk assessments, and an estimation of the efficacy of longevity interventions in animals and humans. While most clocks are trained to predict chronological age, clocks have been developed to predict more complex composite biological age outcomes, at least in humans. These composite outcomes can be made up of a combination of phenotypic data, chronological age, and disease or mortality risk. Here, we develop the first such composite biological age measure for mice: the mouse phenotypic age model (Mouse PhenoAge). This outcome is based on frailty measures, complete blood counts, and mortality risk in a longitudinally assessed cohort of male and female C57BL/6 mice. We then develop clocks to predict Mouse PhenoAge, based on multi-omic models using metabolomic and DNA methylation data. Our models accurately predict Mouse PhenoAge, and residuals of the models are associated with remaining lifespan, even for mice of the same chronological age. These methods offer novel ways to accurately predict mortality in laboratory mice thus reducing the need for lengthy and costly survival studies.

## Introduction

Despite being the same chronological age, some individuals of the same species become frailer, lose function, and die earlier than others. For instance, certain 70-year-old humans have noticeably more skin wrinkling for their age^1^, have acquired cancers and heart disease much earlier, and experience cognitive decline at a greater rate^2^. Conversely, centenarians have been shown to exhibit many traits that are typical of individuals decades younger^3^. In addition to being easily observed, these phenotypic differences reflect heterogeneity at the genomic, transcriptomic, cellular, and tissue levels that can be quantified^4^. The variability in these age-related changes in humans of the same age suggests that the rate of biological aging varies within a species.

One of the most widely-used measures of biological age in recent years are DNA methylation clocks, which assess methylation status at dozens to hundreds of CpG sites to predict chronological age^5,6^. Although the residuals of these clocks (the difference between predicted and actual age) may provide some information about biological age^7^, there is now a focus on the development of biomarkers that measure biological age directly^8^. These are often referred to as “second generation” aging clocks. For example, DNAme PhenoAge was developed from large-scale human studies and uses DNA methylation data to predict PhenoAge, a composite outcome made up of chronological age modulated by health status and mortality risk^9^. PhenoAge comprises 9 clinical blood measurements and chronological age, which are regressed to predict the probability of mortality in the next 10 years and subsequently transformed into an age proxy. DNAme PhenoAge is the DNA methylation-based algorithm that predicts this biological age outcome. Other examples of DNAme-based second-generation aging clocks include DNAme GrimAge^10^, DunedinPoAm^11^, and bAge^12^. These second-generation DNAme biomarkers have been shown to associate with disease risk, mortality, and frailty more strongly than clocks that only predict chronological age (e.g., Horvath’s multi-tissue clock^5^)^13^. In particular, the DNAme PhenoAge clock has been shown to outperform chronological age clocks in predicting mortality, physical performance, and Alzheimer’s disease^9^.

While robust epigenetic clocks that predict chronological age have been developed for numerous species, second-generation aging clocks are only available for humans, largely due to the difficulty and cost in obtaining the paired phenotypic and omic data required to train the models for other species. Model organism second generation clocks would enable more efficient, objective, and clinically relevant outcomes to be measured in pre-clinical testing of diverse longevity interventions before progressing to clinical development. In a previous study^14^, we presented the AFRAID clock that predicts mouse mortality based on items included in the mouse clinical frailty index^15^, a non-invasive assessment of phenotypic measures of age-related decline. This model performs well at stratifying mice by likelihood of death based on a Kaplan-Meier survival analysis, but it lacks molecular data.

Here, we develop novel second-generation clocks for mice based on phenotypic data and blood samples collected in a large cohort of C57BL/6NIA mice across their lifespan. To our knowledge, this is the first study to have matched repeated samples for molecular analysis with complete health and lifespan data. We first develop a mouse phenotypic age model, Mouse PhenoAge, using mouse frailty index data and blood composition measurements from up to five distinct time points, regressed to predict the probability of mortality in the next 4 months, and transformed into an age proxy. Using genome-wide DNAme and metabolomic data generated from mouse blood, we then develop omic based clocks to predict PhenoAge (DNAm PhenoAge, Mtb PhenoAge, and MultiOmic PhenoAge), and show they have a significant correlation with time-to-death across distinct age-groups of mice. Our work promises to accelerate aging research by providing robust second-generation aging clocks that can be used in interventional preclinical research studies.

## Results

### Development of the first mouse phenotypic age (Mouse PhenoAge) model

The development of omic based models of Phenotypic Age involves a three-step process: 1) train and apply a model to predict the remaining lifespan from phenotypic measures, 2) transform the predicted remaining lifespan to an estimated “PhenoAge”, and 3) train a model to predict PhenoAge with omics data^9^. To this end, we conducted a longitudinal analysis of complete blood counts (CBC), 31 frailty measures, and multiomic data from 94 C57BL/6NIA mice at up to 5-time points between mid-life and death, representing 233 total time points (**Figure 1A**). The mice included in this study spanned two cohorts: (1) 45 male and female C57BL/6NIA mice, and (2) 44 male C57BL/6NIA mice.

**Figure 1.**
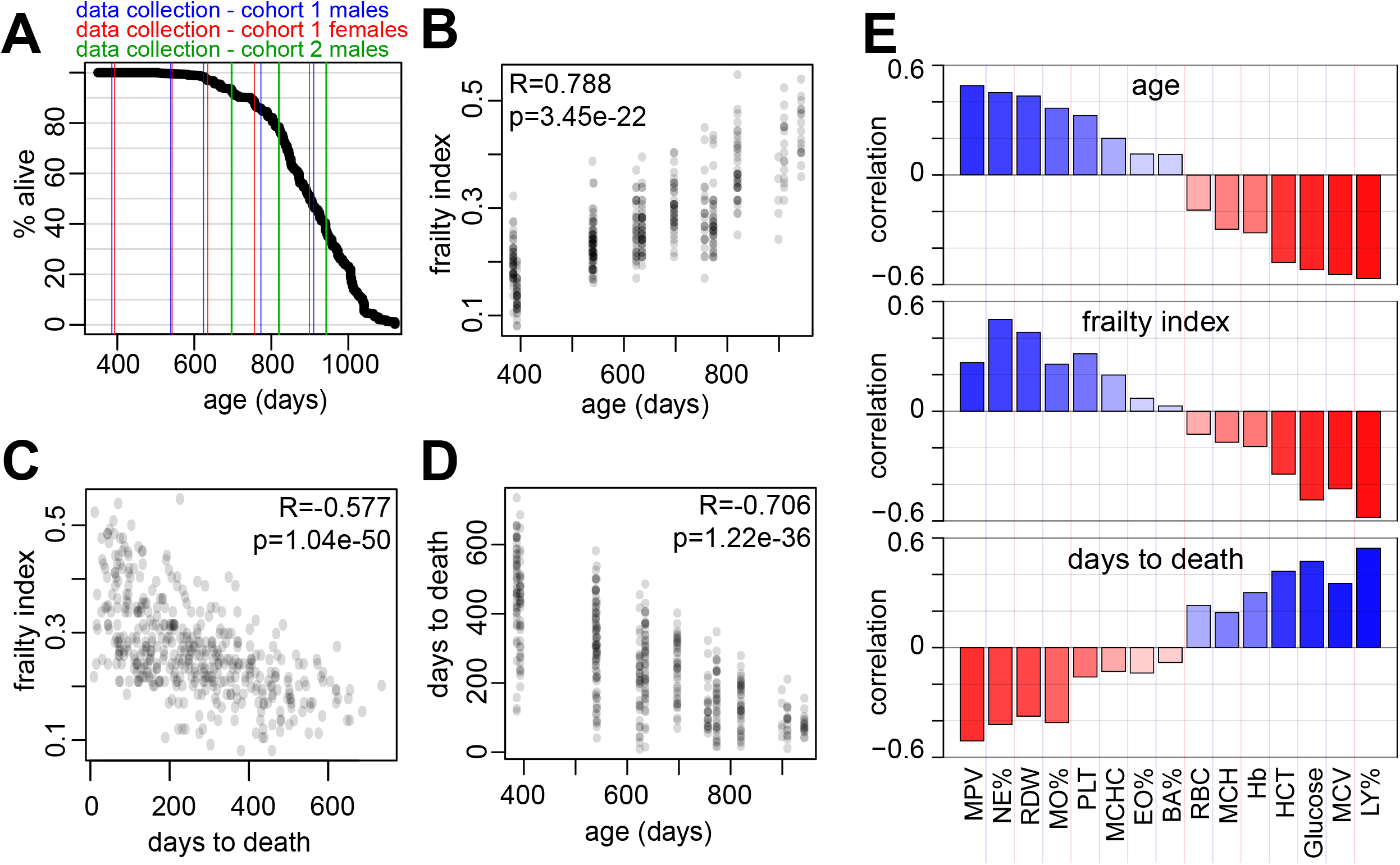
Evaluation of complete blood count (CBC) and frailty index measures and their relationships with primary outcomes. (A) Survival analysis of the 94 C57BL/6NIA mice followed through their life until death. Longitudinal analysis of CBCs, frailty index measures, and multi-omic data were performed for up to 5-time points between mid-life and death (vertical colored lines). (B) Scatter plot of frailty index vs age. (C) Scatter plot of frailty index vs days to death. (D) Scatter plot of days to death vs age. For (B)-(D) pearson correlations (R) and corresponding p values are shown on each graph. (E) Spearman correlations of each CBC measure and age, frailty index, and days to death.

We first assessed the suitability of the phenotypic measures for modeling by evaluating their relationships with chronological age, frailty index, and remaining lifespan (days tio death). As expected, frailty index scores and the individual frailty measures are correlated with age and remaining lifespan, reflecting the physiological manifestation of the aging process (**Figure 1B-C, Supplementary Figure 1, Table S1**). Age is also correlated with remaining lifespan (**Figure 1D**). Additionally, we observed similar associations of multiple CBC measures with age, remaining lifespan, and frailty index (**Figure 1E, Table S1**). Overall, more than 50% of features across the CBC and frailty measures were correlated with remaining lifespan with a correlation coefficient greater than 0.2 (**Table S1**), suggesting sufficient features for multivariate modeling of remaining lifespan.

Using frailty index measures, CBC measures, and chronological age as predictors, we trained an elastic net regression model to predict days to death (Step 1). This model was used to predict the remaining lifespans at each time point for each mouse, which were then transformed into PhenoAge predictions using simple linear equations (detailed in Methods) (Step 2). As expected, PhenoAge was strongly correlated with chronological age (one of the features) and days-to-death (the predicted outcome) (**Figure 2A**, first row, **Table S2**). However, in order for PhenoAge to represent a useful surrogate measure for biological age, differences in PhenoAge among similarly aged mice must explain variability in health outcomes (namely, increased mortality risk). To test this, we calculated PhenoAge acceleration (PhenoAgeAccel, defined by the difference between PhenoAge and chronological age, or PhenoAge-ChronologicalAge) for each sample, and measured its correlation with remaining lifespan at each time point (**Figure 2B**). Positive PhenoAgeAccel values reflect predicted biological age acceleration (higher mortality risk than expected for age), while negative PhenoAgeAccel values reflect predicted biological age deceleration. At each timepoint, PhenoAgeAccel displays negative correlations with remaining lifespan, indicating that mice with greater than expected PhenoAge are at higher risk of death, while mice with lower-than-expected PhenoAge are at lower risk of death (**Figure 2B**, first row). In particular, PhenoAgeAccel measured at later time points were most predictive of mortality (T2-T5), while measurements at the earliest time point were least predictive (T1), possibly due to lower signals differentiating mortality risk so early in life (**Table S3**). When mice are stratified into highest, middle, and lowest PhenoAgeAccel groups (tertiles) for each time point, mice within the highest PhenoAgeAccel tertile tended to live shorter than the other tertiles (**Figure 2C**, first row). Altogether, these findings demonstrate that mice with greater PhenoAge than their chronological age are at higher risk of death, suggesting that PhenoAge is an accurate surrogate measure for biological age that is useful for estimating future mortality risk.

**Figure 2.**
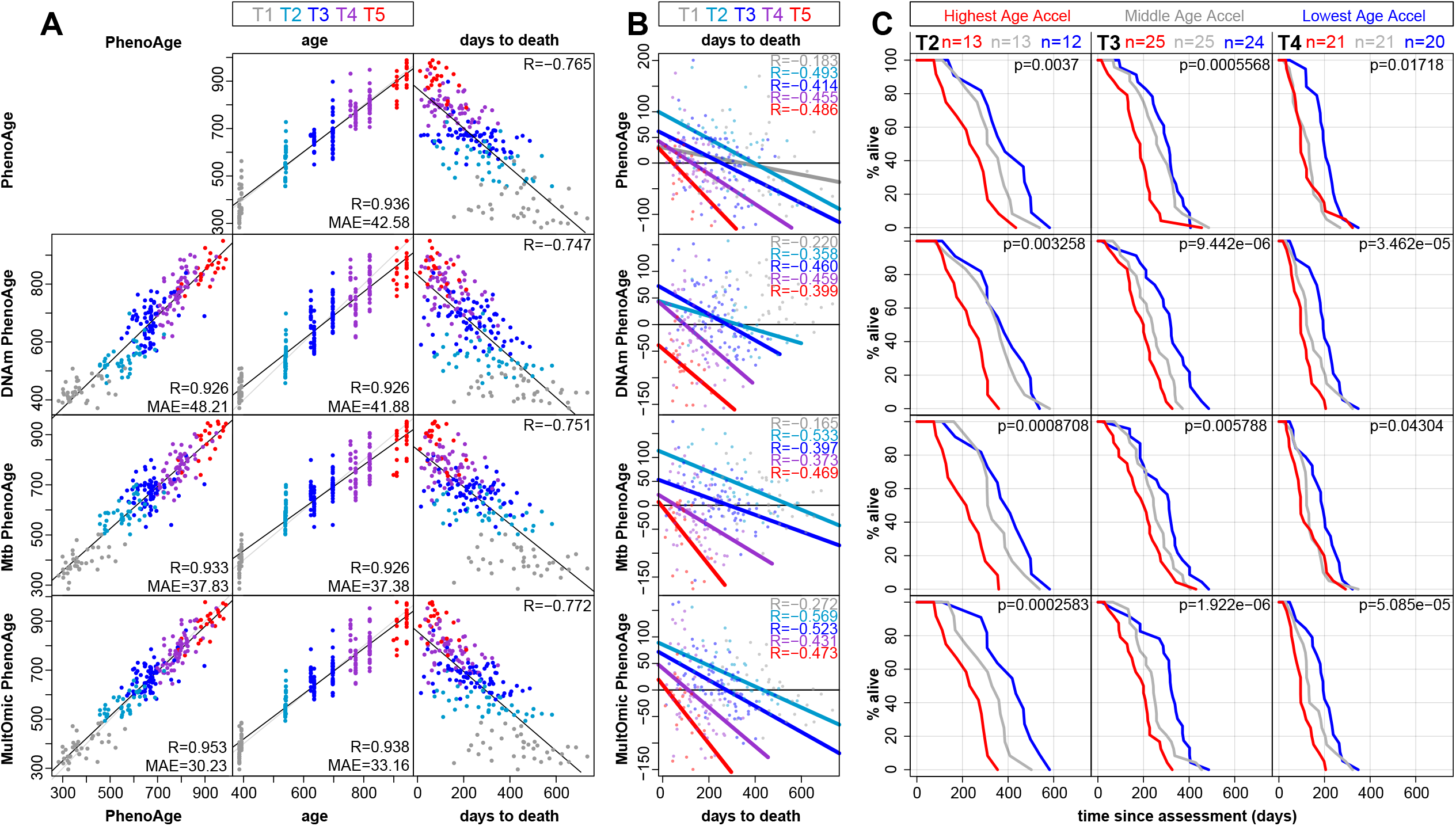
PhenoAge and omic-based PhenoAge models predict mortality risk. (A) Scatter plots of PhenoAge (first row) or predicted Phenoage (second-fourth rows) vs PhenoAge, age, and remaining lifespan. Points are colored by sample time point, and pearson correlations (R) and corresponding median absolute errors (MAE) are shown on each graph. (B) Remaining lifespan was plotted against the Acceleration (age-subtraction) of PhenoAge or predicted PhenoAge. Points, regression lines and spearman correlations are colored by time point. Regression lines are only plotted when p<0.05. (C) For each time point, mice were split into tertiles representing the highest, middle, and lowest PhenoAge (or predicted PhenoAge) acceleration. Percent survival within each group was plotted against time-since-assessment (time from which data was collected on each mouse). P-values represent significance of log-rank test comparing highest and lowest age-accelerated groups. All corresponding statistics for Figure 2 can be found in **Table S2** and **S3**.

### Second generation epigenetic clocks to predict PhenoAge

The PhenoAge model estimates a mouse’s future mortality risk, or biological age, using frailty and CBC measures. However, the collection of these data is cumbersome, time-consuming, and requires special instrumentation. To enable wider utilization of PhenoAge, we sought to construct models to predict PhenoAge using blood-based assays that are readily available through commercial vendors - microarray-based DNA methylation and mass-spectrometry-based metabolomics (Step 3). Such proxy models can simplify the assessment of the health status and future mortality risk of mice at a single point in time from blood.

To begin, we generated DNA methylation data from 233 mouse whole-blood samples collected across nearly all available mice and time points using the Infinium Mouse Methylation BeadChip. We first assessed the suitability of the methylation data for modeling by evaluating the relationship of methylation state at each CpG site (∼285,000) with PhenoAge (**Supplementary Figure 2**). We identified thousands of CpGs whose methylation levels were highly correlated with PhenoAge. with considerably more CpGs that were negatively correlated than positively correlated, possibly reflecting the general trend toward genome-wide hypomethylation that occurs with age^16^.

To train a model to predict PhenoAge with DNAm data (DNAmPhenoAge), we performed elastic net regression with leave-one-mouse-out cross-validation (LOMOCV) to avoid leakage across time points. As expected, out-of-sample DNAmPhenoAge predictions were highly correlated with age and remaining lifespan (features of the mPhenoAge model) (**Figure 2A**, second row). DNAmPhenoAge predictions were also highly correlated with PhenoAge (the target), suggesting DNA methylation data can captures the signal of phenotypic measures utilized in the training of PhenoAge (**Figure 2A**, compare first and second rows). To test whether DNAmPhenoAgeAccel (DNAmPhenoAge - ChronologicalAge) is predictive of mortality risk, we evaluated the correlation between DNAmPhenoAgeAccel and remaining lifespan. As with PhenoAgeAccel, DNAmPhenoAgeAccel was correlated with days to death among 4 of the 5 time points (|R|>0.3, p<0.05, **Figure 2B, Table S2**). When samples within each time point are grouped into highest, middle, and lowest DNAmPhenoAgeAccel groups, mice with the highest DNAmPhenoAgeAccel died significantly sooner than mice with the lowest DNAmPhenoAgeAccel (**Figure 2C,** second row). These data suggest that DNA methylation data can accurately predict the surrogate biological age measure PhenoAge, and can be used to predict age-adjusted mortality risk up to 18 months before death.

### Metabolomic and multiomic clocks to predict PhenoAge

Metabolomics data has been used to predict chronological age in human datasets^17^, but the power of this type of data to predict biological age in preclinical models was unknown. To train a model to predict PhenoAge with metabolomic data (MtbPhenoAge), we performed elastic net regression with LOMOCV as above. MtbPhenoAge displayed high correlations with age, PhenoAge, and remaining lifespan, similar to DNAmPhenoAge (**Figure 2A**, third row). Like PhenoAgeAccel and DNAmPhenoAgeAccel, MtbPhenoAgeAccel was correlated with days to death among most age groups (**Figure 2B**, third row, **Table S2**). Likewise, mice with the highest MtbPhenoAgeAccel died significantly sooner than mice with the lowest MtbPhenoAgeAccel (**Figure 2C**, third row). These data suggest that metabolomic data can accurately predict the biological age measure of PhenoAge, and can be used to accurately predict mortality risk of mice with similar performance to DNA methylation data.

While DNA methylation and metabolomic data can independently predict mortality risk, we questioned whether they would be even more predictive when combined into a single model. To this end, we trained a model to predict PhenoAge with both DNA methylation and metabolomic data (“MultiOmicPhenoAge”).

MultiOmicPhenoAge had a slightly greater correlation with age, PhenoAge, and days to death than PhenoAge, DNAmPhenoAge, or MtbPhenoAge (**Figure 2A**, fourth row). MultiOmicPhenoAgeAccel also better predicted differences in survival at most time points (**Figure 2B,C**).

### PhenoAge-based clocks outperform first generation clocks

To benchmark the performance of our PhenoAge clocks against traditional first generation clocks, we used the same methods to build clocks to predict chronological age (DNAmAge, MtbAge, MultiOmicAge) in the same mouse dataset. DNAmAge was able to accurately predict age, and showed a negative correlation with days to death (**Figure 3A, Table S4**). DNAmAgeAccel (DNAmAge - ChronologicalAge) was associated with remaining lifespan for 2 of the 5 time points (**Figure 3B-C, Table S5**), compared to 4 of the 5 time points for DNAmPhenoAgeAccel. While MtbAge predictions were similarly correlated to chronological age (**Figure 3A**), MtbAgeAccel was not associated with remaining lifespan or differences in survival at any timepoint (**Figure 3B-C**, second row). Out-of-sample predictions from the MultiOmicAge model were the most predictive of chronological age out of every model (R=0.98, **Figure 3A**), however MultiOmicAgeAccel was only associated with remaining lifespan for one time point (**Figure 3B-C**, third row). Altogether, these data suggest that epigenetic, metabolomic, and multiomic first generation clocks are highly predictive of chronological age, but do not predict mortality as well as second generation PhenoAge clocks.

**Figure 3.**
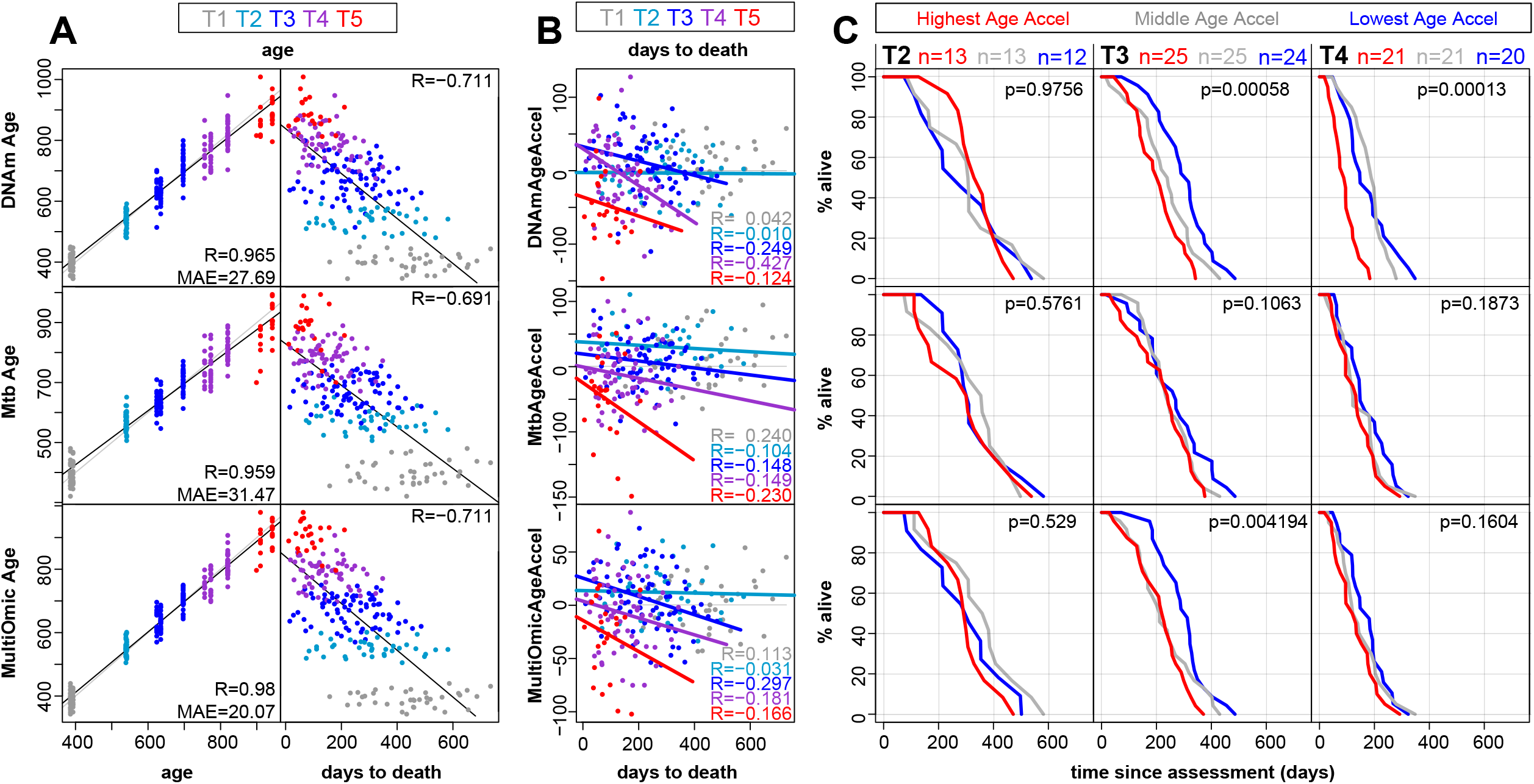
First generation clocks accurately predict age but not mortality risk. (A) Scatter plots of chronological age predictions vs actual age (in days) and remaining lifespan (days to death). Points are colored by sample time point, and pearson correlations (R) and corresponding median absolute errors (MAE) are shown on each graph. (B) The remaining lifespan was plotted against the age acceleration (predicted age minus chronological age). Points, regression lines, and spearman correlations (R) are colored by time point.Regression lines are only plotted when p<0.05. (C) For each time point, mice were split into tertiles representing the highest, middle, and lowest age acceleration. Percent survival within each group was plotted against time-since-assessment (time from which data was collected on each mouse). P-values represent significance of log-rank test comparing the highest and lowest age-accelerated groups. All corresponding statistics for Figure 2 can be found in **Table S4** and **S5**.

## Discussion

Here we present the first second-generation molecular clocks for mice: DNAmPhenoAge, MtbPhenoAge, and MultiOmicPhenoAge. Our findings suggest that blood-based DNA methylation and/or metabolomics data can accurately predict future, age-adjusted mortality risk in mice. These models allow researchers to assess the impact of interventions or diseases on biological age and mortality risk, without the need for lengthy lifespan pre-clinical studies.

For years accurate chronological age clocks have been available for mice based on a variety of technologies and biomolecules^18–21^. However, until now, there were no mouse aging biomarkers that predicted mortality risk directly. In this study, we provide the first mouse phenotypic age models (Mouse PhenoAge), developed using frailty and complete blood count data collected on a large cohort of mice tracked longitudinally until death. Mouse PhenoAge accurately predicts mortality risk, confirming that frailty index and blood composition data capture relevant factors to accurately establish time to death for an individual mouse. Of note, we observed an increasing strength of correlation with remaining lifespan in Mouse PhenoAges of older mice, which should inform the design of studies that plan to employ this tool.

In this study, we also develop the first second-generation aging clocks for mice, using omics to directly predict biological age outcomes. While first-generation DNAm-based chronological age predictors for mice already existed ^22,23^, until now, there were no models to predict other health outcomes or mortality risk. The metabolomic, DNAme, and multi-omic PhenoAge predictions all outperformed first-generation clocks in terms of mortality prediction, highlighting the advantage of second-generation clocks over first. These clocks also out-performed our previously published lifespan estimator, the AFRAID clock^14^, that used only frailty index data as input. However, these approaches should be seen as complementary since the frailty index is inexpensive and easily assessed in aging mice, whereas DNAme BeadChip and/or metabolomic analysis is more costly and time-consuming. In particular, our multiomic model, which is the most accurate model for mortality prediction in mice to date, requires separate readouts from two omics platforms, making it most likely to be used in aging intervention experiments thought to have a moderate effect (i.e., requiring maximum sensitivity) in smaller-scale experiments. Using these relatively expensive and laborious biomarkers to develop equally informative but more targeted and efficient biomarkers^24^ could increase their use and should be the subject of future studies.

Interestingly, although the multiomic predictor was the most accurate for both age and mortality prediction, the integration of more data types did not increase predictive capacity much above the single datasets alone (eg. r=-0.77 vs r-0.75, **Figure 2**). This implies that both the DNA methylation and plasma metabolomics datasets contain enough information about risks of morbidity and mortality to be able to accurately predict their incidence. Further work should examine the specific features associated with aging outcomes in order to understand possible underlying biological determinants.

Mice are the most common mammalian model organism used in aging research. Given their crucial role in preclinical research enabling both the discovery and translation of diverse interventions to the clinic, it is surprising that “second generation” clocks had not already been developed for mice. One of the limiting factors was likely a lack of suitable data sets to train the predictive models. In humans, large cohort studies comprising many researchers across multiple institutions have been collecting the necessary data and biological samples required to train such clocks for decades. In mice, these types of efforts are not as centralized and, as far as we are aware, this is the first study to collect the matched phenotype and molecular data necessary to train omics based biological age predictors. The clocks and data we provide will buoy basic aging research, allowing for multiomic based estimation of biological age and mortality risk in preclinical studies.

The current lack of matched omics and phenotype datasets for mice also means that we were not able to validate our clocks in external datasets, and future studies should validate the applicability and accuracy of these models in other mouse datasets, and/or with interventions. Several other limitations of this study should also be considered. First, we use only mice that are older than 13 months of age, meaning that our models are not expected to work in younger mice. Since mPhenoAge is designed to reflect mortality risk in the 4 months after data collection, the biomarker is not expected to be as predictive in younger animals regardless. Similarly, there were relatively fewer females used in this study, meaning the models and conclusions are biased toward male mice. Additionally, data in this study were collected longitudinally, meaning that values from individual mice are not necessarily independent from collection to collection.

Overall, here we adopt for the first time the ‘PhenoAge’ concept from humans to mice, building models based on frailty and whole blood counts that can accurately predict mortality risk even in mice as young as 18 months of age. We also built the first second-generation molecular clocks for mice, using both DNA methylation and plasma metabolomics data to accurately predict biological age and mortality risk. These models provide important tools for the field to accelerate preclinical aging studies and highlight the diverse data types capable of measuring aging signatures.

## Methods

### Animals

All experiments were conducted according to protocols approved by the Institutional Animal Care and Use Committee (Harvard Medical School). C57BL/6NIA mice were ordered from the National Institute on Aging (NIA, Bethesda, MD) and housed at Harvard Medical School. Mice were housed in rooms with ventilated caging and a 12:12 light cycle, at 71 °F with 45–50% humidity. Mice were tracked in two separate cohorts and fed either LabDiet #5053 or AIN-93G. Mice were group housed to begin (4-5 mice per cage), although over the period of the experiment, mice died and mice were left singly housed. Mice had a normal median (914 days) lifespan, surpassing those cited by Jackson Labs (878 days)^25,26^ and the NIA (793 days) (https://www.nia.nih.gov/research/dab/aged-rodent-colonies-handbook/animalinformation), demonstrating that the mice were maintained and aged in healthy conditions. Mice were only euthanized if determined to be moribund by an experienced researcher or veterinarian based on exhibiting at least two of the following: inability to eat or drink, severe lethargy or persistent recumbence, severe balance or gait disturbance, rapid weight loss (>20%), an ulcerated or bleeding tumor, and dyspnea or cyanosis. In these rare cases, the date of euthanasia was taken as the best estimate of death. For all other euthanized mice, their data was excluded from any analysis of time-to-death. The first cohort tracked was part of an ongoing research experiment to assess the effect of Nicotinamide Mononucleotide (NMN) treatment on healthspan and lifespan^27^. Only untreated mice were considered in the analysis of omic data.

### Mouse Frailty Assessment

Frailty was assessed using the mouse clinical FI^15,28^. Briefly, mice were scored either 0, 0.5 or 1 for the degree of deficit they showed in each of 31 health-related items with 0 representing no deficit, 0.5 representing a mild deficit, and 1 representing a severe deficit. More details can be found here: http://frailtyclocks.sinclairlab.org

### Blood collection and processing

Mice were fasted for 5-6 hours, anaesthetized with isoflurane (3-5%), and then blood was collected from the submandibular vein with a lancet (maximum 10% of mouse body weight, approx. 200-300 ul). One drop of blood was used to measure fasting glucose using a blood glucose monitor and disposable strips, and the rest was collected into a tube containing 20ul of 0.5M EDTA. Blood was mixed and stored on ice. 30ul was removed for whole blood counts with the Hemavet 950 (Drew Scientific). The remaining whole blood was centrifuged at 1500g for 15 mins, plasma was removed and frozen at -80°C for subsequent metabolomics, and the remaining cell pellet for also frozen at -80°C for subsequent DNA extraction.

### Plasma metabolomics

Global metabolomics analysis was completed by Metabolon. Samples were prepared using the automated MicroLab STAR® system (Hamilton Company), and analyzed using Ultrahigh Performance Liquid Chromatography-Tandem Mass Spectroscopy (UPLC-MS/MS).

### DNA methylation

DNA from PBMCs was extracted in two ways. 131 samples were extracted by Autogen using the AutoGenprep 965 and their standard workflow for blood DNA in small volumes. Post extraction these samples were cleaned up with the addition of 10% 20mg/ml Rnase A, 30 minutes incubation at 37°C, SPRI bead wash and isolation, then elution in 10mM Tris-HCl pH8. The remaining 103 samples were extracted in-house. Briefly, red blood cells were removed with RBC lysis buffer (155 mM NH4Cl, 12 mM NaHCO3, 0.1 mM EDTA, pH 7.3), and then the remaining cells were digested overnight at 65°C with proteinase K (20mg/ml) in TESR buffer (10mM Tris-HCl, 25 mM EDTA, 0.5% SDS, 20ug/mL RNAse). DNA was washed and isolated using SPRI beads and eluted in 10 mM Tris-HCl pH8. Concentrations were determined using the Qubit fluorometer (ThermoFisher), normalized, and approx. 500ng was sent to FOXO Technologies for DNA methylation analysis with the Infinium Mouse Methylation BeadChip. Data was processed by FOXO biosciences from the raw input (IDAT) to CpG methylation values for each probe with standard methods detailed previously^23^.

### mPhenoAge modeling

To build the PhenoAge model, we first trained a univariate linear regression model to predict days to death from age. This results in a regression model (DaysToDeath = m * Age + b). Separately, we trained an elastic net regression model to predict days to death with frailty index measures, CBC measures, and chronological age as predictors. To calculate a PhenoAge for each mouse, we take the predicted survival time from the PhenoAge model and transform it into an equivalent age by determining what chronological age would correspond to the same expected number of days to death, based on the relationship established predicting days to death from age alone: (PhenoAge = (PredictedDaysToDeath -b) / m). To train DNAm PhenoAge, Mtb PhenoAge and MultiOmic PhenoAge models, we trained elastic net regression models to predict PhenoAge using DNA methylation, Metabolomic, and DNA methylation + Metabolomic data, respectively. The process of training models and calculating all PhenoAge models were done using LOOCV for each mouse separately to prevent data leakage.

### Metabolomic data pre-processing

We used raw metabolite data (peak area) and only included metabolites with less than 25% missing values. To address batch effects, we calculated the mean metabolite value for the baseline time point across all mice and compared it to the mean value of all other time points. If the mean of the baseline was significantly lower (0.05x, compared to the mean of all other time points) or significantly higher (10x compared to the mean of all other time points), we removed the metabolite from the analysis. We normalized each sample by dividing the metabolite value by the median of metabolite values for that sample to account for any collection batch effects. We log-transformed metabolites for all models except for the time-to-death models. To eliminate outliers, we winsorized all features to the top and bottom 1% and 99% of values for all models except the time-to-death, which we winsorized to 3%. We mean- and standard deviation-normalized all features before running the regression.

### Frailty index transformation for predictive modeling

For modeling (but not for correlations), the data was pre-transformed to reduce noise in scoring phenotypes between time points by eliminating phenotypic regression. So, when a mouse was scored as a full (1) or partial phenotype (0.5), the maximum score was taken in perpetuity. For example, considering the scores for one mouse for one frailty item ‘Tail Stiffening’ at 5 time points: if the original score was [0, .5, 1, .5, 1], these would be transformed to [0, .5, 1, 1, 1]. This transformation reduced error caused by the subjectiveness of some of these ratings because in theory, these individual metrics shouldn’t decrease over time. However, excluding this process didn’t have a substantial effect on the results.

### Predictive modeling from omic data

All modeling analysis was done in Python using the package ‘scikit-learn’ to perform elastic net regression. For each model, we included a binary variable for sex in addition to the omic data. To generate the final models, we combined the data from both lifespan cohorts and used leave-one-out cross-validation (LOOCV) to find the best model parameters and assess model performance. For methylation data, in each fold of the cross-validation, we subset the CpGs down to the 500-1000 CpGs most correlated with the dependent variable (depending on the model and dependent variable).

### Kaplan Meier survival curves and statistical analysis

Kaplan-Meier survival analyses were produced using data on time to death from the beginning of the study for each month supplied to the surv_fit() and ggsurvplot() function from the survminer package (version 0.4.9) in R. To compare timepoints stratified based on the quantile of multi-omic clock predictions, the surv_pvalue() function was used to compute logrank test p-values.

## Contributions

The idea to build multi-omic clocks for mice was initially conceived by A. E. Kane with input from D. A. Sinclair, D. S. Vogel, D. L. Vera, and K. Chwalek. A. E. Kane collected all phenotypic data and processed biosamples used to generate the multi-omic datasets. P. T. Griffin conceived of building a mouse phenotypic age model and performed the analysis to produce mPhenoAge. Data pre-processing and elastic-net regression modeling to predict phenotypes were done by J. Kras, E. Ramos, I. Bishof, A. Butler, and D. L. Vera. P. T. Griffin and D. L. Vera wrote and organized the manuscript with input from A. E. Kane and other authors. All authors reviewed the final manuscript.

## Acknowledgments

We thank the staff of the Harvard Medical Area animal facility for their excellent care of our mouse cohorts.

## References

1. Tsukahara, K. et al. Comparison of age-related changes in wrinkling and sagging of the skin in Caucasian females and in Japanese females. J. Cosmet. Sci. 55, 351–71 (2004).

2. Jaul, E. & Barron, J. Age-Related Diseases and Clinical and Public Health Implications for the 85 Years Old and Over Population. Front. Public Heal. 5, 335 (2017).

3. Shaffer, J. Centenarians, Supercentenarians: We Must Develop New Measurements Suitable for Our Oldest Old. Front. Psychol. 12, 655497 (2021).

4. López-Otín, C., Blasco, M. A., Partridge, L., Serrano, M. & Kroemer, G. Hallmarks of aging: An expanding universe. Cell doi:10.1016/j.cell.2022.11.001.

5. Horvath, S. DNA methylation age of human tissues and cell types. Genome Biol 14, R115–R115 (2013).

6. Hannum, G. et al. Genome-wide Methylation Profiles Reveal Quantitative Views of Human Aging Rates. Mol. Cell 49, 359–367 (2013).

7. Chen, B. H. et al. DNA methylation-based measures of biological age: meta-analysis predicting time to death. Aging Albany Ny 8, 1844–1859 (2016).

8. Bell, C. G. et al. DNA methylation aging clocks: challenges and recommendations. Genome Biol 20, 249 (2019).

9. Levine, M. E. et al. An epigenetic biomarker of aging for lifespan and healthspan. Aging Albany Ny 10, 573–591 (2018).

10. Lu, A. T. et al. DNA methylation GrimAge strongly predicts lifespan and healthspan. Aging Albany Ny 11, 303–327 (2019).

11. Belsky, D. W. et al. Quantification of the pace of biological aging in humans through a blood test, the DunedinPoAm DNA methylation algorithm. Elife 9, e54870 (2020).

12. Bernabeu, E. et al. Refining epigenetic prediction of chronological and biological age. Genome Med 15, 12 (2023).

13. Horvath, S. & Raj, K. DNA methylation-based biomarkers and the epigenetic clock theory of ageing. Nat. Rev. Genet. 19, 371–384 (2018).

14. Schultz, M. B. et al. Age and life expectancy clocks based on machine learning analysis of mouse frailty. Nat Commun 11, 4618 (2020).

15. Whitehead, J. C. et al. A Clinical Frailty Index in Aging Mice: Comparisons With Frailty Index Data in Humans. J. Gerontol. Ser. A: Biomed. Sci. Méd. Sci. 69, 621–632 (2014).

16. Day, K. et al. Differential DNA methylation with age displays both common and dynamic features across human tissues that are influenced by CpG landscape. Genome Biol. 14, R102 (2013).

17. Robinson, O. et al. Determinants of accelerated metabolomic and epigenetic aging in a UK cohort. Aging Cell 19, e13149 (2020).

18. Horvath, S. DNA methylation age of human tissues and cell types. Genome Biol 14, R115–R115 (2013).

19. Unfried, M. et al. LipidClock: A Lipid-Based Predictor of Biological Age. Front. Aging 3, 828239 (2022).

20. Meyer, D. H. & Schumacher, B. BiT age: A transcriptome_Jbased aging clock near the theoretical limit of accuracy. Aging Cell 20, e13320 (2021).

21. Tanaka, T. et al. Plasma proteomic signature of age in healthy humans. Aging Cell 17, e12799 (2018).

22. Mozhui, K. et al. Genetic loci and metabolic states associated with murine epigenetic aging. Elife 11, (2022).

23. Zhou, W. et al. DNA methylation dynamics and dysregulation delineated by high-throughput profiling in the mouse. Cell Genom. 2, 100144 (2022).

24. Griffin, P. T. et al. TIME-Seq reduces time and cost of DNA methylation measurement for epigenetic clock construction. Nature Aging, (2024) 10.1038/s43587-023-00555-2 PMID: 38200273

25. Festing, M. F. W. Inbred Strains in Biomedical Research. (1979) 10.1007/978-1-349-03816-9.

26. Kunstyr I, Leuenberger H. G. Gerontological data of C57BL/6J mice. I. Sex differences in survival curves. J Gerontol. 30(2):157–62. doi: 10.1093/geronj/30.2.157

27. Kane A. E et al. Long-term NMN treatment increases lifespan and healthspan in mice in a sex dependent manner. BioRxiv. 2024. 10.1101/2024.06.21.599604 PMID: 38979132

28. Kane, A. E., Ayaz, O., Ghimire, A., Feridooni, H. A. & Howlett, S. E. Implementation of the mouse frailty index1. Can. J. Physiol. Pharmacol. 95, 1149–1155 (2017).

