## Supplementary Figures and Tables for "Multiomic clocks to predict phenotypic age in mice"

### Supplemental figures

**Supplementary Figure 1.** Correlations of the individual frailty measures with age and remaining lifespan.

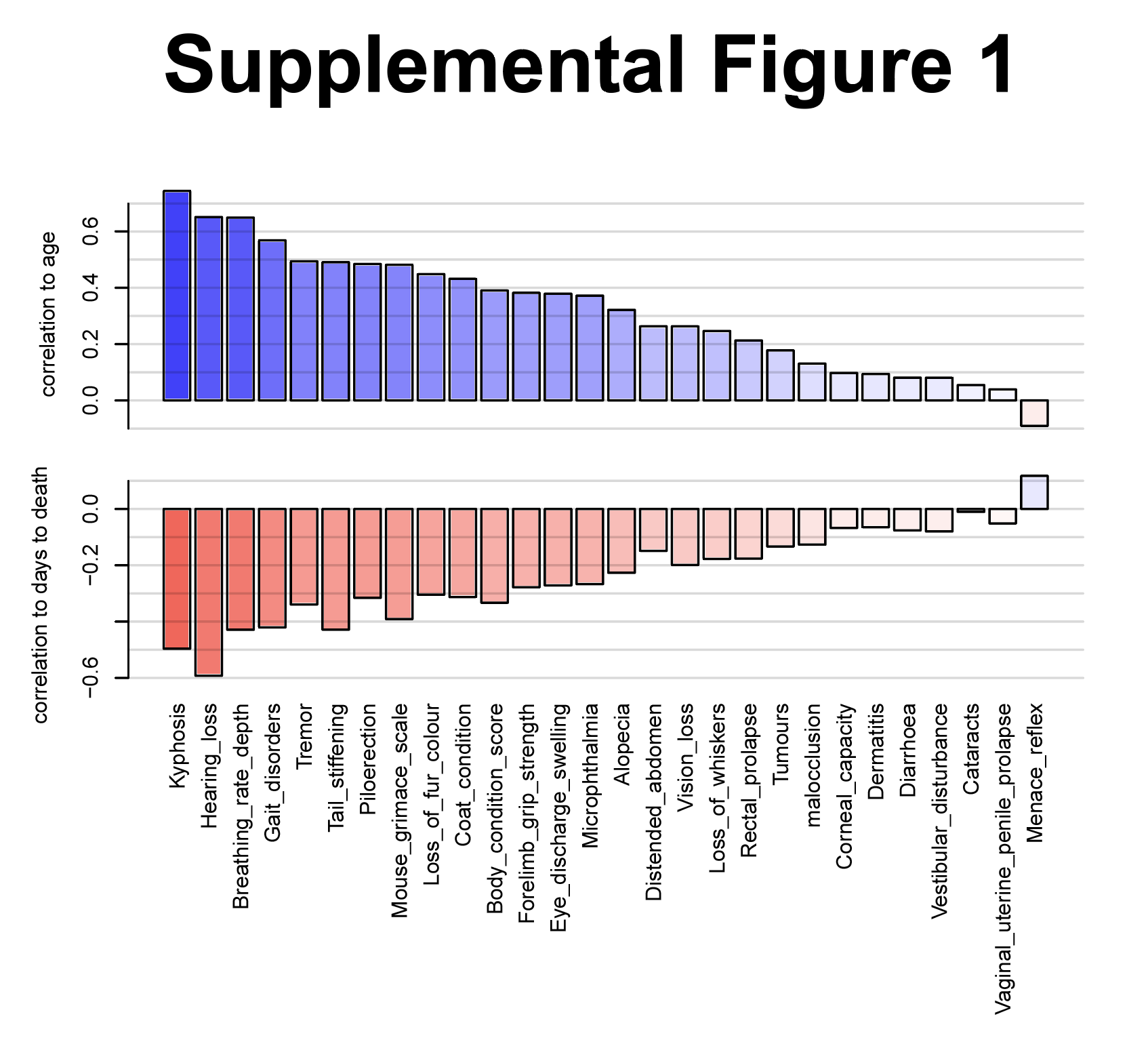

**Supplementary Figure 2.** Correlation of methylation levels at each measured CpG with PhenoAge.

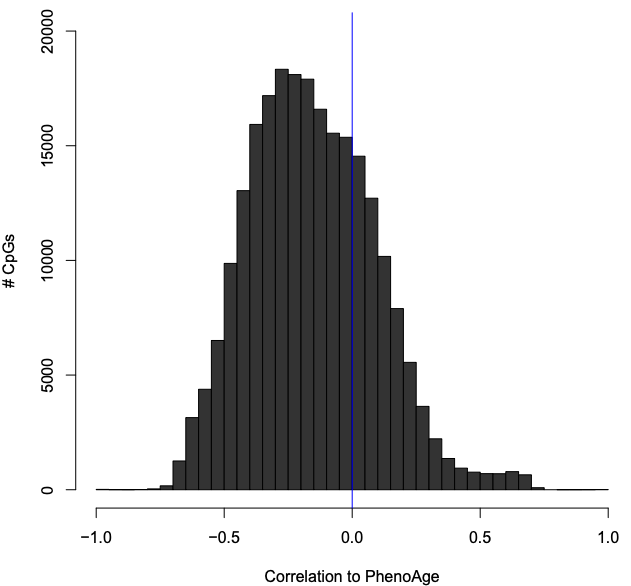

### Supplemental tables

**Table S1.** Correlations of CBC and frailty index measures with Age, Frailty Index, and Remaining Lifespan.

| **Measure** | **Age**  **(Correlation coefficient)** | **Age**  **(P Value)** | **FI**  **(Correlation coefficient)** | **FI**  **(P Value)** | **DaysToDeath**  **(Correlation coefficient)** | **DaysToDeath**  **(P Value)** |
| --- | --- | --- | --- | --- | --- | --- |
| **Body weight** | 0.101437436 | 0.121771909 | 0.174492102 | 0.007462261 | 0.055891873 | 0.394731422 |
| **Glucose** | -0.55526331 | 5.97484E-19 | -0.50355441 | 2.36579E-15 | 0.474270039 | 1.43407E-13 |
| **NE%** | 0.549536447 | 3.99168E-18 | 0.564745015 | 2.94347E-19 | -0.474981453 | 2.49916E-13 |
| **LY%** | -0.591454443 | 2.15431E-21 | -0.583355749 | 1.0037E-20 | 0.552157231 | 2.57118E-18 |
| **MO%** | 0.27490063 | 4.96228E-05 | 0.20607078 | 0.002568739 | -0.370196347 | 2.74507E-08 |
| **EO%** | 0.119717091 | 0.082023987 | 0.088131986 | 0.201206126 | -0.185827581 | 0.006660314 |
| **BA%** | 0.096531136 | 0.161370393 | 0.050331555 | 0.46602128 | -0.109865987 | 0.11070163 |
| **RBC** | -0.279157133 | 3.74265E-05 | -0.173693658 | 0.011297758 | 0.267496246 | 8.01713E-05 |
| **Hb** | -0.352868157 | 1.39714E-07 | -0.215069303 | 0.001675926 | 0.280444884 | 3.58469E-05 |
| **HCT** | -0.565721945 | 2.47814E-19 | -0.386952717 | 5.55967E-09 | 0.433244777 | 4.12375E-11 |
| **MCV** | -0.537540252 | 2.84802E-17 | -0.396126913 | 2.231E-09 | 0.304785289 | 6.1986E-06 |
| **MCH** | -0.174853232 | 0.010755968 | -0.082291538 | 0.232820971 | 0.047267721 | 0.493632883 |
| **MCHC** | 0.268070536 | 7.72813E-05 | 0.225172647 | 0.000961064 | -0.18888851 | 0.005800031 |
| **RDW** | 0.447670996 | 7.63206E-12 | 0.459466598 | 1.80911E-12 | -0.412298194 | 4.16269E-10 |
| **PLT** | 0.343824091 | 3.02842E-07 | 0.303529564 | 7.1493E-06 | -0.135641063 | 0.049102652 |
| **MPV** | 0.466898893 | 7.09663E-13 | 0.309671826 | 4.31287E-06 | -0.464124958 | 1.00899E-12 |
| **Alopecia** | 0.330882425 | 2.2058E-07 | 0.587816802 | 3.8847E-23 | -0.247055846 | 0.000134348 |
| **Loss of fur colour** | 0.500867541 | 2.89986E-16 | 0.570177266 | 1.41437E-21 | -0.354562248 | 2.44765E-08 |
| **Dermatitis** | 0.070207665 | 0.285869155 | 0.202959076 | 0.001846654 | -0.01951129 | 0.767036586 |
| **Loss of whiskers** | 0.283190464 | 1.13528E-05 | 0.483391868 | 4.78223E-15 | -0.188827044 | 0.003817173 |
| **Coat condition** | 0.410532852 | 6.28673E-11 | 0.620563253 | 2.65736E-26 | -0.2753137 | 1.93918E-05 |
| **Tumours** | 0.205302641 | 0.001591774 | 0.342748085 | 7.49961E-08 | -0.148822439 | 0.022783264 |
| **Distended abdomen** | 0.243171563 | 0.000172354 | 0.270428916 | 2.75283E-05 | -0.10964544 | 0.094264377 |
| **Kyphosis** | 0.782148562 | 1.44668E-49 | 0.766319987 | 1.79132E-46 | -0.507559427 | 1.00213E-16 |
| **Tail stiffening** | 0.529887964 | 2.43102E-18 | 0.602415572 | 1.67246E-24 | -0.428976047 | 6.81108E-12 |
| **Gait disorders** | 0.567832282 | 2.24447E-21 | 0.710929537 | 2.4915E-37 | -0.437690265 | 2.27196E-12 |
| **Tremor** | 0.448319854 | 5.70246E-13 | 0.473349931 | 1.80881E-14 | -0.302479026 | 2.43937E-06 |
| **Forelimb grip strength** | 0.378417881 | 2.20949E-09 | 0.443628705 | 1.0558E-12 | -0.294353976 | 4.63813E-06 |
| **Body condition score** | 0.38455974 | 1.1518E-09 | 0.576721115 | 3.82267E-22 | -0.287227438 | 8.02027E-06 |
| **Vestibular disturbance** | 0.128100042 | 0.050332078 | 0.194218263 | 0.002849537 | -0.134827902 | 0.039317832 |
| **Hearing loss** | 0.616802391 | 6.41002E-26 | 0.480008508 | 6.88173E-15 | -0.577193864 | 3.47396E-22 |
| **Cataracts** | 0.064809692 | 0.32358045 | 0.190544835 | 0.003433264 | -0.007457963 | 0.909653552 |
| **Corneal capacity** | 0.107784728 | 0.100020357 | 0.212179436 | 0.001092314 | -0.067998434 | 0.300293032 |
| **Eye discharge swelling** | 0.361789437 | 1.20585E-08 | 0.465162887 | 5.76906E-14 | -0.251793503 | 9.86055E-05 |
| **Microphthalmia** | 0.354501112 | 2.46217E-08 | 0.483124834 | 4.34633E-15 | -0.258932174 | 6.11673E-05 |
| **Vision loss** | 0.21912072 | 0.000737989 | 0.310351755 | 1.28447E-06 | -0.134920233 | 0.039182104 |
| **Menace reflex** | -0.078559065 | 0.231252632 | -0.007338617 | 0.911093315 | 0.141136737 | 0.030909936 |
| **malocclusion** | 0.131705015 | 0.04601862 | 0.151598446 | 0.02145466 | -0.128501418 | 0.051621233 |
| **Rectal prolapse** | 0.232246291 | 0.000339966 | 0.326989113 | 3.11205E-07 | -0.199452493 | 0.002172639 |
| **Vaginal uterine penile prolapse** | 0.114600639 | 0.080216172 | 0.085570004 | 0.192113112 | -0.050923375 | 0.438156758 |
| **Diarrhoea** | 0.1013321 | 0.122161645 | 0.17297394 | 0.008005401 | -0.100130193 | 0.12667748 |
| **Breathing rate depth** | 0.657984241 | 2.08348E-30 | 0.675968393 | 1.3477E-32 | -0.44172239 | 1.35247E-12 |
| **Mouse grimace scale** | 0.47116889 | 2.47114E-14 | 0.485765235 | 2.93382E-15 | -0.356097574 | 2.10899E-08 |
| **Piloerection** | 0.57212196 | 9.61674E-22 | 0.581804198 | 1.35567E-22 | -0.367079874 | 7.10121E-09 |

**Table S2.** Summary statistics from Figure 2A - Median Absolute Error (MAE) and Pearson Correlations (including p-values) between predicted and actual PhenoAge.

|  | PhenoAge | Age | Days to Death |
| --- | --- | --- | --- |
| PhenoAge |  | MAE = 42.58 R = 0.9364 p = 1.8195e-107 | R = -0.7646 p = 3.802e-46 |
| DNAm PhenoAge | MAE = 48.21 R = 0.9262 p = 2.9574e-100 | MAE = 41.88 R = 0.9264 p = 2.1787e-100 | R = -0.7473 p = 4.6064e-43 |
| Mtb PhenoAge | MAE = 37.83 R = 0.9334 p = 3.2213e-105 | MAE = 37.38 R = 0.9262 p = 2.8048e-100 | R = -0.7516 p = 8.2841e-44 |
| MultiOmic PhenoAge | MAE = 30.23 R = 0.9532 p = 1.8254e-122 | MAE = 33.16 R = 0.9383 p = 5.5715e-109 | R = -0.7724 p = 1.251e-47 |

**Table S3.** Summary statistics from Figure 2B – Spearman correlation between predicted Age Acceleration and remaining lifespan at each assessed time point.

|  | T1 | T2 | T3 | T4 | T5 |
| --- | --- | --- | --- | --- | --- |
| PhenoAgeAccel | R = -0.1832 p = 0.2778 | R = -0.4927 p = 0.001674 | R = -0.414 p = 0.0002219 | R = -0.4553 p = 0.0002007 | R = -0.486 p = 0.02185 |
| DNAm PhenoAgeAccel | R = -0.2201 p = 0.1906 | R = -0.3584 p = 0.02716 | R = -0.4603 p = 3.25e-05 | R = -0.459 p = 0.000175 | R = -0.3985 p = 0.06624 |
| Mtb PhenoAgeAccel | R = -0.1647 p = 0.3301 | R = -0.5333 p = 0.0005662 | R = -0.3976 p = 0.0004117 | R = -0.373 p = 0.00283 | R = -0.4694 p = 0.02754 |
| MultiOmic PhenoAgeAccel | R = -0.2716 p = 0.1039 | R = -0.5695 p = 0.0001901 | R = -0.5234 p = 1.445e-06 | R = -0.4314 p = 0.0004636 | R = -0.4725 p = 0.02636 |

**Table S4.** Summary statistics from Figure 3A - Median Absolute Error (MAE) and Pearson Correlations (including p-values) between predicted and actual Age.

|  | Age | Days to Death |
| --- | --- | --- |
| DNAm Age | MAE = 27.69 R = 0.9652 p = 3.6868e-137 | R = -0.7107 p = 2.7228e-37 |
| Mtb Age | MAE = 31.47 R = 0.9591 p = 4.1834e-129 | R = -0.6905 p = 1.7514e-34 |
| MultiOmic Age | MAE = 20.07 R = 0.9803 p = 1.9569e-165 | R = -0.7109 p = 2.5275e-37 |

**Table S5.** Summary statistics from Figure 3B – Spearman correlation between predicted Age Acceleration and remaining lifespan at each assessed time point.

|  | T1 | T2 | T3 | T4 | T5 |
| --- | --- | --- | --- | --- | --- |
| DNAm AgeAccel | R = 0.04163 p = 0.8067 | R = -0.01062 p = 0.9496 | R = -0.2487 p = 0.03144 | R = -0.4267 p = 0.0005431 | R = -0.1242 p = 0.5818 |
| Mtb AgeAccel | R = 0.2399 p = 0.1527 | R = -0.1035 p = 0.5365 | R = -0.1482 p = 0.2046 | R = -0.1485 p = 0.2495 | R = -0.2302 p = 0.3027 |
| MultiOmic AgeAccel | R = 0.113 p = 0.5054 | R = -0.03149 p = 0.8512 | R = -0.2973 p = 0.009594 | R = -0.1809 p = 0.1595 | R = -0.1666 p = 0.4588 |
